# Cell size regulation in budding yeast does not depend on linear accumulation of Whi5

**DOI:** 10.1101/2020.01.20.912832

**Authors:** Felix Barber, Ariel Amir, Andrew W. Murray

**Affiliations:** Department of Molecular and Cellular Biology, Harvard University, Cambridge, MA 02138, USA; School of Engineering and Applied Sciences, Harvard University, Cambridge, MA 02138, USA; FAS Center for Systems Biology, Harvard University, Cambridge, MA 02138, USA

**Keywords:** Whi5, Start, cell size control, budding yeast, single-cell time-lapse microscopy

## Abstract

Cells must couple cell cycle progress to their growth rate to restrict the spread of cell sizes present throughout a population. Linear, rather than exponential, accumulation of Whi5, was proposed to provide this coordination by causing a higher Whi5 concentration in cells born at smaller size. We tested this model using the inducible *GAL1* promoter to make the Whi5 concentration independent of cell size. At an expression level that equalizes the mean cell size with that of wild-type cells, the size distributions of cells with galactose-induced Whi5 expression and wild-type cells are indistinguishable. Fluorescence microscopy confirms that the endogenous and *GAL1* promoters produce different relationships between Whi5 concentration and cell volume without diminishing size control in the G1 phase. We also expressed Cln3 from the GAL1 promoter, finding that the spread in cell sizes for an asynchronous population is unaffected by this perturbation. Our findings contradict the previously proposed model for cell size control in budding yeast and demonstrate the need for a molecular mechanism that explains how cell size controls passage through Start.

**Author Contributions:** FB performed the experiments, data analysis and simulations. All authors designed the experiments and wrote the manuscript.

**Significance Statement:** Despite decades of research, the question of how single cells regulate their size remains unclear. Here we demonstrate that a widely supported molecular model for the fundamental origin of size control in budding yeast is inconsistent with a set of experiments testing the model’s key prediction. We therefore conclude that the problem of cell size control in budding yeast remains unsolved. This work highlights the need for rigorous testing of future models of size control in order to make progress on this fundamental question.

## Introduction

Cells from all domains of life display size control (1–4), coupling their cell size and cell cycle progression to reduce the variability in size observed throughout a population. We refer readers to the following recent reviews for further discussion (5–8). Despite this widespread behavior, the parameters that cells monitor as proxies for their size and the mechanisms by which these parameters control progress through the cell cycle remain unclear. In the budding yeast *S. cerevisiae*, studies of the cell cycle have produced several hypotheses for the mechanism of size control (9–14). However, no consensus has yet been reached that favors one model above all others. Several models include a role for Whi5, a transcriptional inhibitor that delays progress through Start (15–17), the transition that commits yeast cells to replicating their DNA and then dividing. Here we test how cell size regulation depends on the expression dynamics of Whi5.

Budding yeast divides asymmetrically, with size control acting in the first cell cycle of the smaller, newborn daughter cells. During their first cell cycle, small daughters have a longer G1 phase, which delays Start relative to the cell cycle of larger daughter cells (3, 18, 19). In contrast, the timing of the budded portion of the cell cycle is roughly independent of cell size (18, 20). The core regulatory network that controls passage through Start has been studied extensively and is outlined in Figure 1 (15, 16, 21–28).

**Figure 1:**
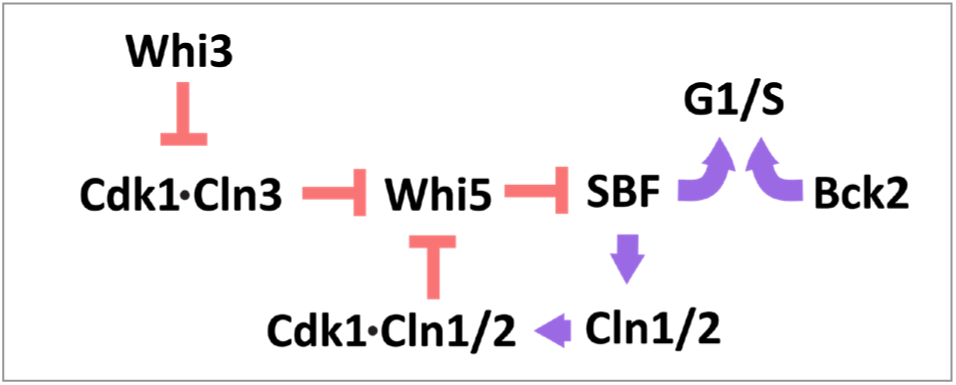
Genetic regulation of passage through Start.

At the end of mitosis, Whi5 enters the nuclei of both daughter and mother cells, where it remains for much of the subsequent G1 phase (15). Its nuclear localization allows Whi5 to bind to and inhibit the heterodimeric transcription factor SBF, preventing it from inducing the expression of genes required for passage through Start (15, 16). The G1 cyclin Cln3 forms a heterodimer with the cyclin-dependent kinase Cdk1, which phosphorylates both Whi5 and SBF at multiple sites, causing Whi5 to unbind from SBF (25, 15, 16, 29, 30). SBF and another transcription factor, MBF, induce genes that drive passage through Start including two other G1 cyclins, Cln1 and Cln2. The expression of Cln1 and Cln2 completes a positive feedback loop that commits cells to passage through Start and leads to the nuclear export of phosphorylated Whi5 (31, 32). The differential lengthening of G1 in small daughter cells occurs prior to Whi5’s nuclear exit, demonstrating that cell size must influence the rate of activation of this positive feedback loop (33).

The genetic circuit that regulates Start also includes Bck2, a protein responsible for Cdk1-indpendent activation of genes that promote passage through Start (27, 34–36). The parallel Bck2- and Cln3-dependent induction of gene expression is revealed by the G1 arrest of *cln3Δ bck2Δ* cells, a behavior not seen in either single mutant (27, 35).

A range of different hypotheses for how cell size could regulate this genetic network have been proposed. We highlight two paradigms of size control: inhibitor dilution or activator accumulation (37, 38). Recent observations support an inhibitor dilution model enacted through Whi5, wherein the growth-mediated dilution of Whi5 relative to a roughly constant concentration of Cln3 during G1 increases the rate of passage through Start as cells grow larger (12, 39). Since the concentration of Whi5 is negatively correlated with daughter cell volume at birth, this dilution mechanism could explain the longer G1 of small daughter cells; smaller cells would need to grow proportionally more to dilute their higher initial concentration of Whi5. This negative correlation between Whi5 concentration at birth and cell volume was proposed to originate from a volume-independent synthesis rate of Whi5 during the budded portion of the cell cycle, with only a small fraction of Whi5 synthesis occurring during the G1 phase. This contrasts with the synthesis rate of Cln3 which, like most other proteins, scales with cell volume. Here we focus on the predictions of the Whi5 dilution model.

## Results

### Perturbing the dynamics of Whi5 expression does not alter the cell size distribution

We tested whether the details of how Whi5 accumulates determine the success of cell size regulation. The negative correlation between Whi5 concentration at birth and cell volume is essential for the Whi5 dilution model to regulate cell size (12). We perturbed this correlation by expressing Whi5 proportionally to cell volume, decoupling Whi5 concentration at birth from cell volume at birth. Our approach also eliminates the periodic nature of Whi5 synthesis, meaning that a synthesis rate proportional to volume would cause Whi5 concentration to reach a constant value, independent of cell size. Since the rate of passage through Start is proposed to decrease with increasing Whi5 concentration, cells would delay Start significantly if the steady state Whi5 concentration was higher or than that of newborn wild-type (WT) daughters, and the perturbed cells would become larger with each cell cycle. Conversely if the perturbed Whi5 level was lower than that of WT cells, the cells with altered Whi5 expression would become progressively smaller. This behavior is illustrated in Figure S1, with simulations validating these predictions (see SI Methods). However, these extremes are not seen experimentally; cells lacking Whi5 are indeed smaller than WT cells, but these cells still display size control, rather than the steadily decreasing cell size that the simple, Whi5-dependent model predicts (15, 16, 40, 41).

We perturbed Whi5 synthesis by placing the *WHI5* gene under the control of the *GAL1* promoter (*P*_*GAL1*_*-WHI5*). This promoter has previously been used to show that overexpressing Whi5 in an otherwise WT background increases mean cell size (15, 42). It is, however, difficult to achieve finely graded control of Whi5 expression in these cells: expression from the *GAL1* promoter is bi-stable at intermediate galactose concentrations and cells metabolize galactose, altering its concentration throughout an experiment. Deleting *GAL1* blocks galactose phosphorylation and metabolism and placing *GAL3* under the control of a strong promoter (*P*_*ACT1*_*-GAL3*) makes expression from *P*_*GAL1*_ vary smoothly with the concentration of galactose in the medium (42). To quantify Whi5 expression, we generated a fusion to Whi5 using the fast-maturing fluorescent protein mVenNB (43). Figure S2 demonstrates that overexpressing Whi5 by exposing these *P*_*GAL1*_-*WHI5*-*mVenNB* cells to high levels of exogenously added galactose generates cells that are larger than *P*_*WHI5*_*-WHI5-mVenNB* cells. For brevity, these cell types will henceforth be abbreviated to *P*_*GAL1*_*-WHI5* and *P*_*WHI5*_*-WHI5*. Figure S2 further shows that, at a high galactose concentration, these *P*_*GAL1*_*-WHI5* cells also maintain a concentration of Whi5 in G1 that is several times larger than that of *P*_*WHI5*_*-WHI5* cells (measured by fluorescence microscopy using fluorescence intensity as a proxy for protein concentration). Despite this, these large cells still maintain a reproducible characteristic cell size. These findings are consistent with previous observations (15, 16), but are inconsistent with the inhibitor dilution model’s prediction that these unusually high levels of Whi5 expression will lead to an unconstrained increase in average cell size over time (see Figure S1).

Since the predictions of an inhibitor dilution model were not validated by overexpressing Whi5, we decided to characterize the cell size control displayed by our *P*_*GAL1*_*-WHI5* cells in greater detail to assess the effect of decoupling Whi5 concentration at birth from cell volume at birth. Tuning the exogenous galactose concentration to 125μM generated cells whose average size was identical to that of *P*_*WHI5*_*-WHI5* cells (Figure S3). Synthesizing Whi5 proportionally to cell volume should eliminate size control during G1 by removing the negative correlation between cell size and Whi5 concentration at birth. If the rate of passage through Start couples to cell size through the Whi5 concentration, as has been claimed (12), the rate of passage through Start will be uncorrelated with cell volume. Because the primary size control in budding yeast occurs during G1, the loss of this control is predicted to generate a substantial broadening of the cell size distribution. Indeed, in the absence of compensatory mechanisms, a constant Whi5 concentration is expected to generate arbitrarily broad distributions of cell size, as demonstrated through simulations in Figure S1 (G-I). Figure S3 shows that in bulk cultures we observed no such broadening of the cell size distribution, inconsistent with the predictions of the Whi5 inhibitor dilution model. Because the Whi5 dilution model was initially tested in cells lacking Bck2, we also compared the size of *P*_*WHI5*_*-WHI5, bck2Δ* and *P*_*GAL1*_*-WHI5, bck2Δ* cells at 125 μM galactose (12). As expected, *bck2Δ* cells were larger than *BCK2* cells, but neither the mean cell size nor the standard deviation in cell size was significantly different between the *P*_*WHI5*_*-WHI5, bck2Δ* and *P*_*GAL1*_*-WHI5, bck2Δ* cells. Aside from these measurements on asynchronous populations of cells (Figure S3), our data was obtained in *BCK2* cells.

To study the distributions of cell size at specific points in the cell cycle we used a microfluidic device to track the growth of immobilized cells in a flow chamber when exposed to 125 μM galactose. We extracted information on 3581 individual cell cycles using a custom-designed algorithm (Methods). Figure 2 shows the average cell size and coefficient of variation (CV = standard deviation / mean) in cell size measured at birth, at Start and at division for daughter and mother cells separately. These observations show a statistically significant, though minor, increase in the average and CV of volume at birth for *P*_*GAL1*_*-WHI5* daughter cells relative to *P*_*WHI5*_*-WHI5* cells. We did not observe statistically significant differences in CV in cell volume between *P*_*GAL1*_*-WHI5* cells and *P*_*WHI5*_*-WHI5* cells at Start or at division. Figure S3 shows cell size statistics for asynchronous populations, measured both using a coulter counter and by imaging large numbers of cells from exponentially growing cultures at a single time-point. Fluorescence imaging in the second approach allowed us to focus on cells in G1 based on the localization of fluorescently tagged Whi5: unbudded cells with Whi5 in the nucleus have not passed Start, whereas those with Whi5 in the cytoplasm have passed Start. However, we were unable to distinguish between mother and daughter cells using this approach. Our measurements on G1 cells using single time-point microscopy and over the full cell cycle using a coulter counter showed a spread in cell size that was independent of whether Whi5 was expressed from the *P*_*GAL1*_ or *P*_*WHI5*_ promoter; both populations had a similar standard deviation and CV. We note, however, that the “smearing” effect of measuring cell size in an asynchronous population may obscure any small increase in the CV for volume at birth. Thus, the only observable difference in cell size between our *P*_*GAL1*_*-WHI5* cells and our *P*_*WHI5*_*-WHI5* cells is a modest increase in the CV in volume at birth in newborn daughter cells. Notably, this finding is inconsistent with the substantially broader cell size distribution predicted by the Whi5 dilution model (see Figure S1).

**Figure 2:**
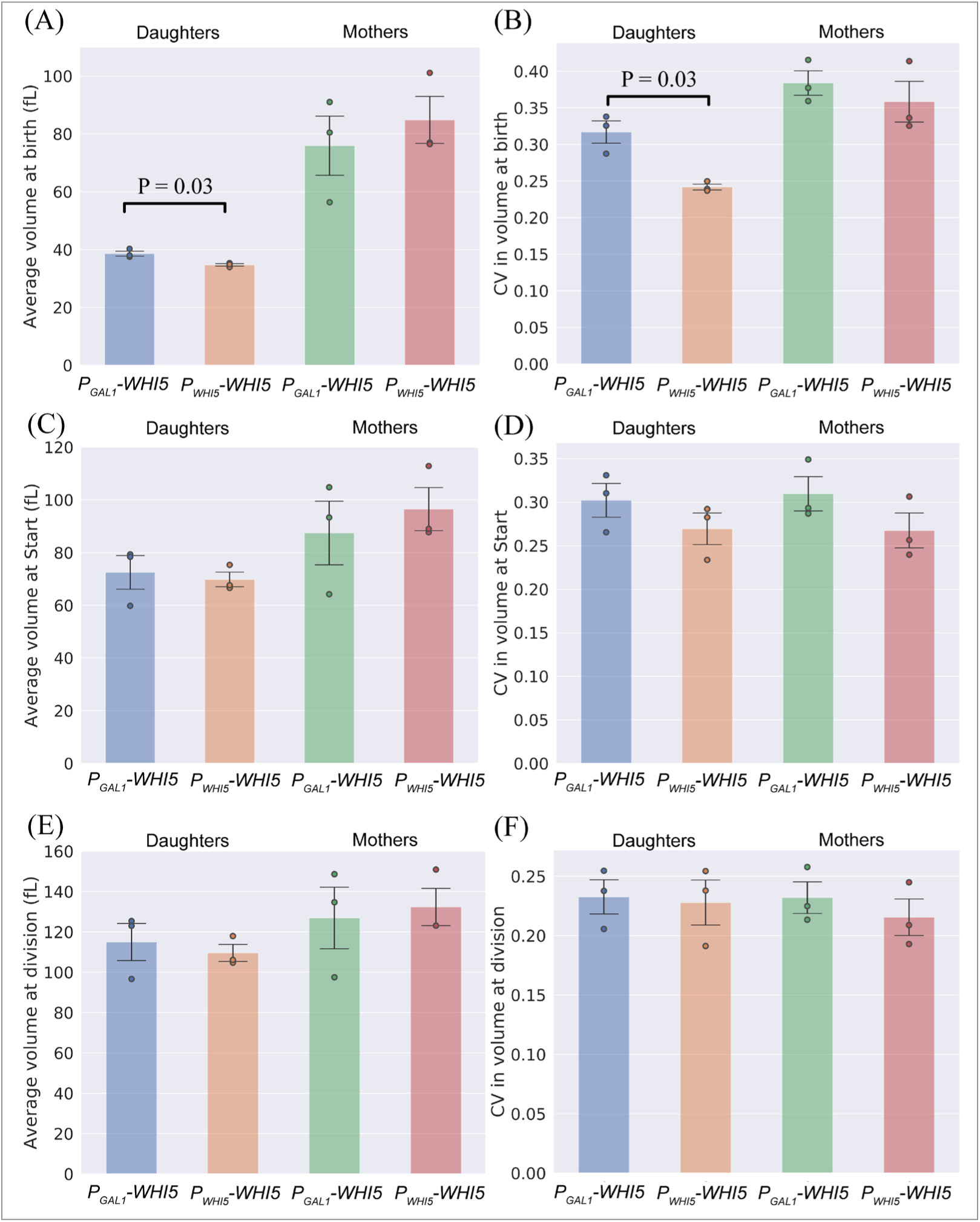
Perturbing Whi5 expression using a galactose-inducible promoter has a minimal effect on the spread in cell size, generating a modest increase in CV at birth, which is not observable at Start or cell division. Cell size distributions are compared for *P*_*GAL1*_*-WHI5* and *P*_*WHI5*_*-WHI5* cell types for volume at birth, at Start, and at division. Volumes were measured via time-lapse microscopy and plotted by cell type (inducible and non-inducible, mothers and daughters). (A) Average size at birth. (B) CV in size at birth. (C) Average size at Start. (D) CV in size at Start. (E) Average size at division. (F) CV in size at division. Values represent the mean across 3 biological replicates of the relevant statistic (average or CV). Measurements taken over a total of 3581 cell cycles: 347 *P*_*GAL1*_*-WHI5* Daughters; 853 *P*_*GAL1*_*-WHI5* Mothers; 800 *P*_*WHI5*_*-WHI5* Daughters; 1581 *P*_*WHI5*_*-WHI5* Mothers. Error bars represent the standard deviation taken across 3 biological replicates. Black lines correspond to statistically significant differences with two-tailed P values less than 0.05 quoted, calculated by comparing daughters and mothers separately between cell types using a Welch’s t-test across biological replicates. Dots correspond to values for individual biological replicates.

Another potential difference between the *P*_*GAL1*_ promoter and the *P*_*WHI5*_ promoter is in the stochasticity of gene expression. Figure S2 demonstrates based on time-point microscopy that the CV in the concentration of Whi5-mVenNB within *P*_*GAL1*_*-WHI5* cells is greater than that of *P*_*WHI5*_*-WHI5* cells. In contrast, our measurements of the CV in cell size at distinct points in the cell cycle, shown in Figure 1, show little difference between our *P*_*GAL1*_*-WHI5* and *P*_*WHI5*_*-WHI5* strains. This observation indicates that any difference in the noisiness of gene expression between these promoters does not lead to a notably broader size distribution. We note that both the magnitude and relative size of these CVs at birth, Start and division are comparable to previous measurements on cells in similar growth conditions (19). If Whi5 concentration is a crucial element in sensing cell size, then greater variability in expression might be expected to lead to a broader size distribution than that seen in WT cells. This contrasts with the unchanged CVs in cell size that we observe and is consistent with our conclusion that the dynamics of Whi5 expression are unimportant in coupling size to passage through Start.

### The dynamics of Whi5 accumulation depend on the promoter driving its expression

We used fluorescence microscopy to confirm that the scaling of Whi5 synthesis with cell volume was perturbed in *P*_*GAL1*_*-WHI5* cells. In the Whi5 dilution model, a linear accumulation of Whi5 leads to a negative correlation between Whi5 concentration and cell volume in newly born cells (12), which was proposed to be essential for controlling the distribution of cell sizes. We therefore measured the correlation between these two quantities in our strains with time-lapse microscopy, using average fluorescence intensity as a proxy for the concentration of Whi5-mVenNB protein at cell birth at the Galactose concentration (125 μM) studied in Figure 2. For *P*_*WHI5*_*-WHI5* cells, we confirmed the previously observed negative correlation between Whi5 concentration at birth and cell volume, as shown in Figure 3B for daughter cells. Figure 3C shows the concentration profile for the expression of *P*_*ACT1*_*-mCherry* in the same cells, with this red fluorescent protein driven by the promoter of the gene *ACT1*. Unlike Whi5-mVenNB, the concentration of mCherry does not decline in larger cells. The *P*_*GAL1*_*-WHI5* cells showed a different pattern of Whi5 accumulation, with no statistically significant correlation between Whi5 concentration and cell volume at birth. Comparing Figures 3B and 3C demonstrates that inducing Whi5 synthesis with the *GAL1* promoter brings the correlation between Whi5 concentration at birth and cell volume at birth closer to that observed for our *P*_*ACT1*_*-mCherry* construct. Because individual time-lapse measurements showed greater variability in their correlations and contained fewer cells we performed our statistical tests on time-lapse data aggregated from at least 3 experimental replicates. Figure S4 shows that *P*_*WHI5*_*-WHI5* cells recapitulate the previously observed decrease in Whi5 concentration throughout G1 (12), whereas the Whi5 concentration remains roughly constant throughout G1 in *P*_*GAL1*_*-WHI5* cells. Figure S5 shows that the average level of both Whi5-mVenNB and mCherry are indistinguishable between cells that express Whi5 from the *WHI5* or *GAL1* promoters.

**Figure 3:**
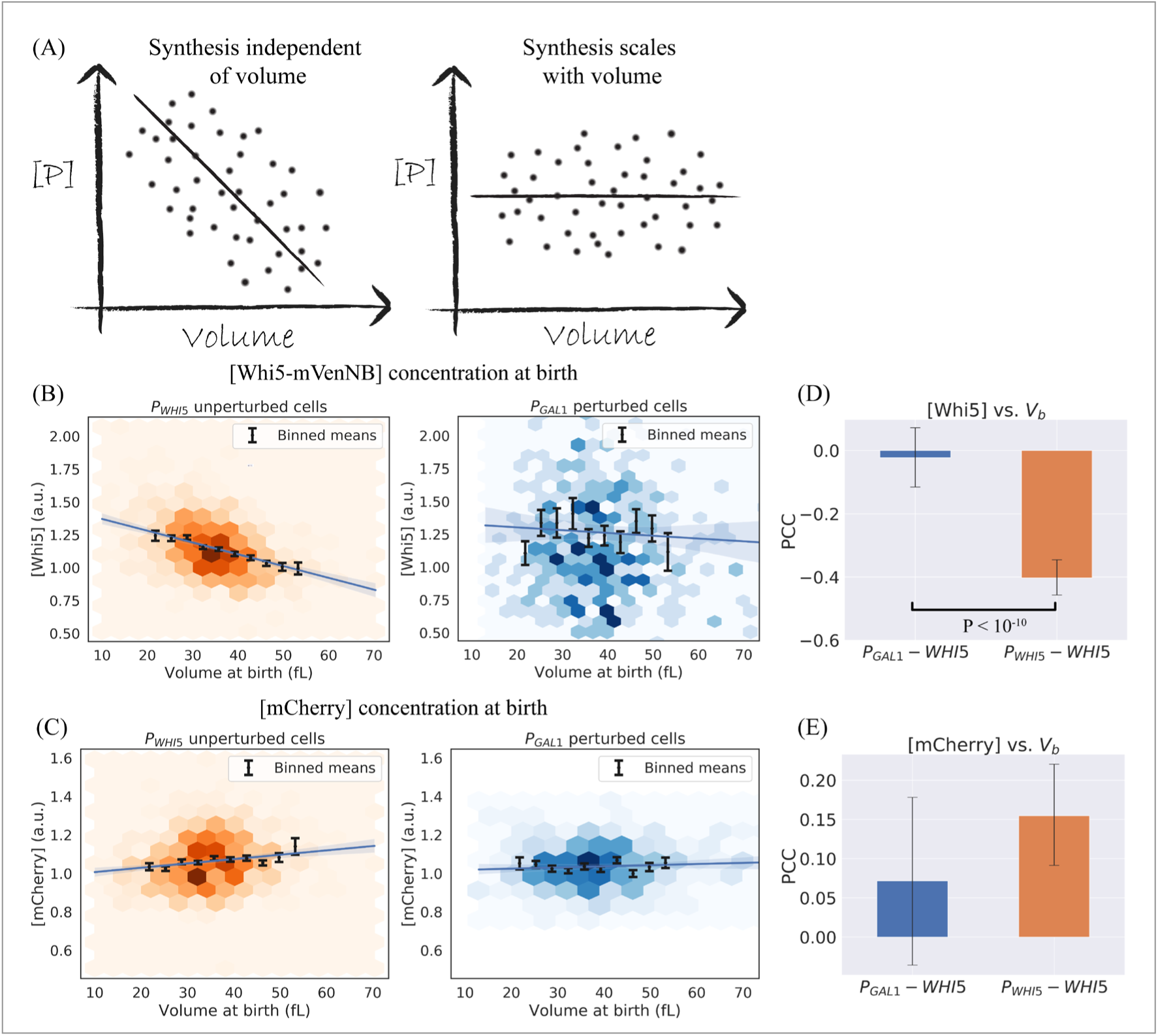
Expressing Whi5 from the *GAL1* promoter alters the relationship between Whi5 concentration and cell size. (A) Illustrations of the predicted correlation of protein concentration with cell size for a gene whose production rate scales linearly with cell volume (50), contrasted with a gene whose synthesis does not scale with cell volume. (B-C) Concentration of fluorescent proteins at cell birth vs volume at birth (*V*_*b*_), grouped by cell type (*P*_*WHI5*_*-WHI5* “unperturbed” cells, and *P*_*GAL1*_*-WHI5* “perturbed” cells) and derived from time-lapse experiments to monitor cell growth. The fluorescence intensity averaged over the cell is used as a proxy for protein concentration. Colored hexagons represent a 2D histogram of datapoints, with darker hexagons showing increased local density of data points. Black lines correspond to averages of the same data binned with respect to *V*_*b*_, with error bars showing the standard error of the mean. Blue lines correspond to linear regression fits, with 95% confidence intervals. Data is compiled from three experiments for each cell type. (B) Whi5 signal. *P*_*WHI5*_*-WHI5* cells (orange) show a negative correlation between Whi5 concentration at birth and cell volume at birth. *P*_*GAL1*_*-WHI5* cells (blue) lose this negative correlation, consistent with Whi5 synthesis being proportional to cell volume. (C) pACT1-mCherry signal. Daughter cells display a weak positive correlation between [mCherry] and cell volume at birth. This correlation is unknown in origin, though it is consistent between *P*_*GAL1*_*-WHI5* and *P*_*WHI5*_*-WHI5* cell types. (D-E) Pearson correlation coefficients (PCC) measured for the datasets plotted in (B) and (C). Error bars correspond to 95% confidence intervals inferred by bootstrapping analysis. Black lines correspond to statistically significant differences with P values less than 0.05 quoted, calculated using a Fisher’s z-transformation on both datasets. (D) PCC values for [Whi5] measured at birth vs. *V*_*b*_ for daughter cells show a statistically significant difference between cell types with P<10^−10^. (E) PCC values for [mCherry] measured at birth vs. *V*_*b*_ for daughter cells shows no statistically significant difference between the two cell types.

Because time-lapse microscopy can alter cell cycle timing and fluorescent proteins show photobleaching after repeated illumination, we also measured the correlation of fluorescence intensity with cell volume for G1 cells in asynchronous cultures. Figure S6 confirms our observations from time-lapse microscopy: the concentration of Whi5-mVenNB expressed from *P*_*WHI5*_ falls with increasing cell volume, whereas the concentration of Whi5-mVenNB expressed from *P*_*GAL1*_ rises with cell volume. This rise is qualitatively similar to the positive correlation observed for mCherry expressed from *P*_*ACT1*_. We conclude that expressing Whi5 from *P*_*GAL1*_ strongly perturbs the correlation between the volume of G1 cells and their Whi5 concentration relative to cells expressing Whi5 from its own promoter, bringing it closer to the behavior observed for expression from the *P*_*ACT1*_ promoter. These findings, combined with those of previous sections, show that the distribution of cell size is unaffected by the difference between a linear and exponential accumulation of Whi5 over time, invalidating a key prediction of the Whi5 dilution model.

### Correlations in cell cycle timing and cell volume

Perturbing the dynamics of Whi5 synthesis failed to change the cell size distribution in asynchronous populations, and only showed a modest increase in CV in volume at birth. To study the effects of this perturbation on size control in greater detail, we examined the dependence of cell cycle timing on cell volume in *P*_*GAL1*_*-WHI5* cells. We defined time the between birth and Start as the duration of Whi5 nuclear localization, while the rest of the cell cycle was defined as the period when Whi5 was excluded from the nucleus. This metric for G1 duration has been validated in previous studies of cell size regulation (33). Figure S7 compares the time spent in the distinct portions of the cell cycle for *P*_*GAL1*_*-WHI5* and *P*_*WHI5*_*-WHI5* cells. The average durations of the entire cell cycle and the pre- and post-Start intervals are statistically indistinguishable between the two populations. Figure 4A shows that the negative correlation between G1 timing and cell volume is seen in both *P*_*GAL1*_*-WHI5* and *P*_*WHI5*_*-WHI5* cells, with *P*_*GAL1*_*-WHI5* cells showing a slightly stronger negative correlation (Fisher z-transformation). The correlation between the duration of the budded portion of the cell cycle (*t*_*budded*_)and cell volume at Start has a slightly stronger negative correlation for *P*_*GAL1*_*-WHI5* cells than for *P*_*WHI5*_*-WHI5* cells (Fisher z-transformation). This finding is consistent with weak size control acting in the budded portion of the cell cycle, but since size control during G1 remains functional in our perturbed cell type, the interpretation of this difference is unclear (Figure 4B). Further, there is no statistically significant difference in the correlation between cell volume at birth and division (a common indicator for the mode of cell size control (8)), when compared between the two cell types. In order to determine whether these differences in the correlations of cell cycle timing with cell volume translated to effects on cell volume, we studied the correlations between the cell volume added in different phases of the cell cycle and cell volume at the start of those phases. Figure S8 shows these correlations for volume added in G1 as a function of volume at birth, volume added during budding as a function of volume at Start, and volume added over the full cell cycle as a function of volume at birth. These datasets show no statistically significant differences between *P*_*GAL1*_*-WHI5* and *P*_*WHI5*_*-WHI5* cells, consistent with size control being conserved between these cell types.

**Figure 4:**
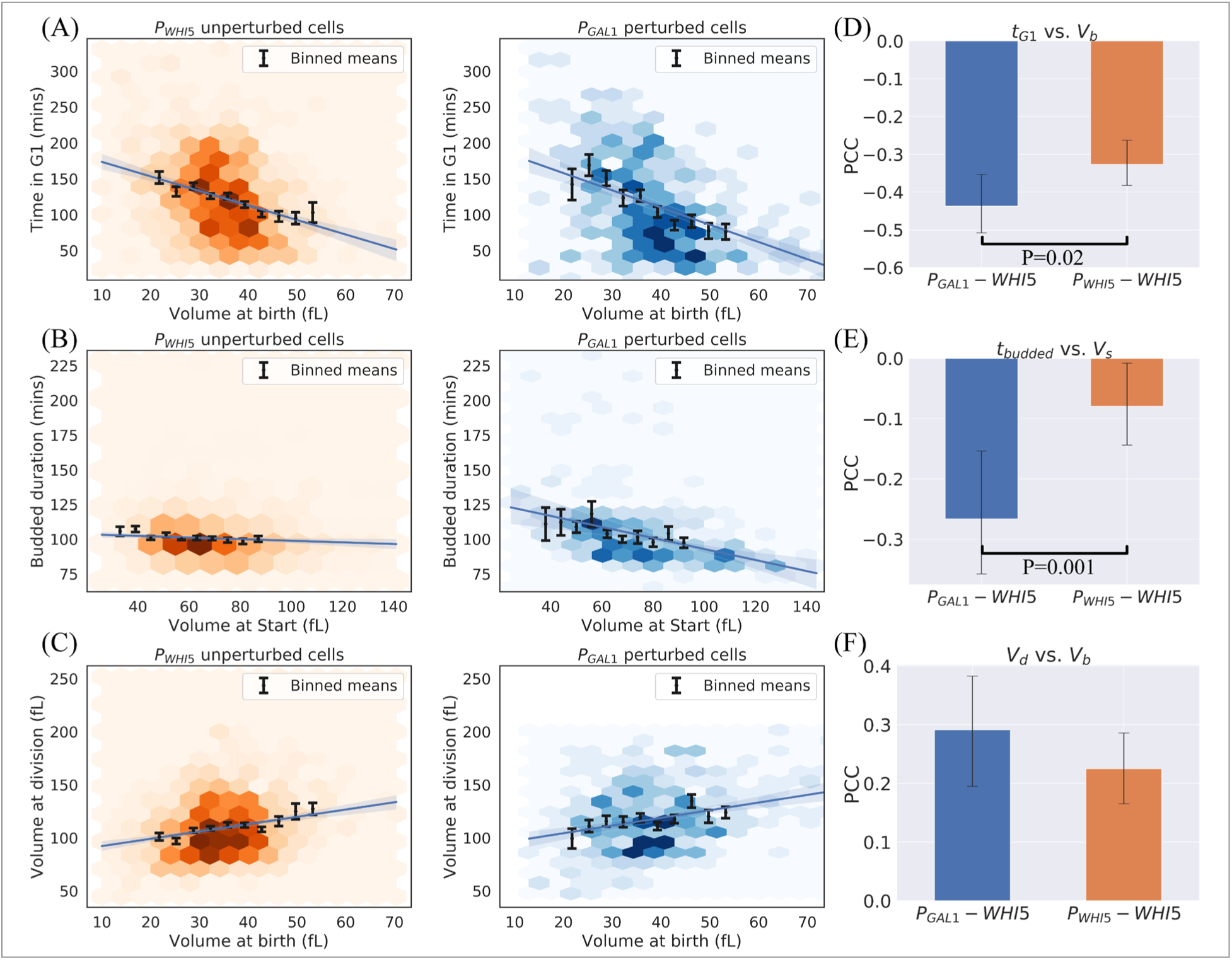
*P*_*GAL1*_*-WHI5* cells retain size control during the G1 phase, in addition to weak size control in the budded portion of the cell cycle. (A-C) Cell cycle correlations for daughter cells. See Figure 3 for details on plotting features for (A-C). Data is compiled from three experiments for each cell type. (A) Plot of time spent in G1 phase (determined by nuclear localization of Whi5) vs. cell volume at birth (*V*_*b*_) for *P*_*GAL1*_*-WHI5* and *P*_*WHI5*_*-WHI5* cells. (B) Plot of time spent in the budded phases (the sum of S-phase, G2, and mitosis, determined by nuclear exclusion of Whi5) vs. cell volume at Start (*V*_*s*_) for *P*_*GAL1*_*-WHI5* and *P*_*WHI5*_*-WHI5* cells. (C) Plot of *V*_*b*_ vs. volume at division (*V*_*d*_). (D-F) PCCs measured for the datasets plotted in (A-C). Error bars show 95% confidence intervals inferred by bootstrapping analysis. Black lines correspond to statistically significant differences with P values less than 0.05 quoted, calculated using a Fisher’s z-transformation on both datasets. (D) PCC values for G1 duration vs. *V*_*b*_ for daughter cells show a statistically significant difference between cell types (P=0.02). This difference is consistent with stronger size control occurring during the G1 phase for *P*_*GAL1*_*-WHI5* cells, not weaker as would be predicted by the inhibitor dilution model. (E) PCC values for budded duration measured at birth vs. *V*_*b*_ for daughter cells shows a statistically significant difference between the two cell types with P=0.001. This difference corresponds to the presence of weak size control during the budded portion of the cell cycle. (F) PCC values for *V*_*b*_ vs. *V*_*d*_ for daughter cells shows no statistically significant difference between the two cell types.

Finally, we tested the dependence of G1 duration on Whi5 concentration at birth. Figure S9 demonstrates that the duration of G1 retains a positive correlation with Whi5 concentration at birth in both *P*_*GAL1*_*-WHI5* and *P*_*WHI5*_*-WHI5* cells, with no statistically significant difference between the correlation coefficients measured for these two cell types. This observation is consistent with our findings and those of others that overexpressing Whi5 leads to an increase in cell size by delaying Start (12, 15, 16).

### Cell size control does not depend on the dynamics of Cln3 expression

Previous studies have focused on the roles of Cln3, as an activator, and Whi5, as an inhibitor, in coupling passage through Start to cell size control (9, 11, 12, 44, 45). To explore whether the details of Cln3 accumulation influence cell size control, we constructed strains where the endogenous copy of *CLN3* was replaced by a galactose-inducible version. As expected, increasing Cln3 expression by increasing the galactose concentration decreased cell size, in contrast to the increased cell size caused by increasing Whi5 expression. Tuning the galactose concentration to 200μM yielded an average cell size corresponding to that of WT cells. In this condition, our *P*_*GAL1*_*-CLN3* cells show no statistically significant difference in mean, standard deviation or CV of their size from that of WT cells (Figure S10). This finding indicates that at the level of an asynchronous population, perturbing the details of Cln3 synthesis does not lead to any observable broadening of the cell size distribution.

## Discussion

We investigated the role of Whi5 in controlling cell size in the budding yeast, *Saccharomyces cerevisiae*. Like previous groups, we find that overexpressing Whi5 makes cells bigger and preventing its expression makes them smaller. Titrating the expression of Whi5 from the galactose-inducible *GAL1* promoter produces populations of cells whose mean size and spread in cell size are indistinguishable from those of cells that express Whi5 from its endogenous promoter. Because expressing Whi5 from *P*_*GAL1*_ makes the rate of Whi5 synthesis scale linearly, rather than sub-linearly, with cell volume, this result is inconsistent with the model that the sublinear scaling of Whi5 synthesis with cell volume plays a critical role in controlling the distribution of cell size. Minor differences in the CV in volume at birth were observed in time-lapse microscopy, however, this modest increase in CV for *P*_*GAL1*_*-WHI5* cells remains inconsistent with the perturbations predicted by an inhibitor dilution model. To confirm that cell size was still being controlled by the length of the interval between cell birth and Start we verified that this timing retains its strong negative correlation with cell volume at birth when Whi5 is expressed from *P*_*GAL1*_. Additionally, whichever promoter Whi5 is expressed from, we see a positive correlation between Whi5 concentration and the interval between birth and Start within a population of cells.

Our results show that the concentration of Whi5 influences the size at which the cell population passes through Start but the dynamics of Whi5 accumulation do not, indicating that Whi5 dilution is unlikely to constrain the spread in the cell size distribution. Our results invalidate a key prediction of the Whi5 dilution model and reveal that any size control model based solely on the dilution of the inhibitor Whi5 is necessarily incomplete. Our results further demonstrate that any effect arising from the anticorrelation between Whi5 concentration and cell volume is not the dominant means by which size control occurs at passage through Start. We note that even if the Whi5 dilution model is not the principal way of controlling the cell size distribution, this mechanism could play a redundant role whose individual effect was too small to be detected within our experiments.

The observations that led to the Whi5 dilution model were made on cells grown in 2% glycerol plus 1% ethanol as a carbon source in order to generate small daughter cells with strong size control. This growth medium differs from the 2% Raffinose carbon source used here. Although both are non-fermentable carbon sources that generate a small average cell size, we cannot rule out the possibility that this difference in carbon source has contributed to the difference between the conclusions of the two studies. However, both studies observed a correlation between Whi5 concentration and G1 phase duration in WT cells and we reproduced the negative correlation between Whi5 concentration and cell volume at birth in G1 that lies at the heart of the inhibitor dilution model. Additionally, Table S2 shows that the duration of cell cycle phases measured in our growth medium are comparable to those published previously, displaying an extended G1 phase but very similar interdivision times (33). Dorsey *et al.* (13) used microscopy to infer absolute protein copy number as cells passed through the cell cycle. This work did not observe any reduction of the Whi5 concentration during G1, after controlling for photobleaching effects, which contrasts with our own and earlier work (12). However, their measurements were acquired using glucose or glycerol as carbon sources. Another recent study proposed that this discrepancy may arise from differences in growth media rather than from photobleaching effects, since their observations showed Whi5 dilution in the G1 phase to be more pronounced for cells grown in media with longer doubling times (44). Our single time-point microscopy experiments for cells in G1 demonstrate that correlation between birth cell volume and Whi5 concentration (Figure 3) that we used as a basis for invalidating the Whi5 dilution model are not a result of photobleaching. We cannot, however, rigorously exclude the effects of photobleaching in our recapitulation of Whi5 dilution during the G1 phase (Figure S4) (12).

Are there viable alternatives to the Whi5 dilution model? A variety of other models have been proposed. They propose the accumulation, throughout G1, of a component that is limiting for passage through Start, such as Cln3 or the SBF subunit Swi4 (9, 13), or the integration of Cln3 activity throughout G1 through its ability to phosphorylate and inhibit Whi5 (11). Attempts to measure Cln3 levels have been limited by its rapid degradation, causing authors to use stabilized Cln3 mutants rather than WT Cln3 (11, 12). To address this concern, a recent study used a self-cleaving linker to allow a fluorescent protein to report on Cln3 translation without affecting Cln3 function or accumulation (45). These authors observed that WT cells experience a pulse of overall protein synthesis in late G1, leading to an increase in Cln3 concentration which drives cells through Start. How this burst couples passage through Start to cell size remains unclear. Cln3 translation is hindered by its long 5’ UTR (44), an effect which may be mediated by the binding of Whi3 (an inhibitor of Cln3 translation) to key motifs in Cln3 mRNA transcripts (47). Collectively, these observations have led to the suggestion that the translation of Cln3 may play a role in sensing cell size at the G1/S transition, for example by having the rate of Cln3 translation increase non-linearly with cell volume during G1 (41). Our measurements on asynchronous populations do not support such a model. Figure S10 demonstrates that replacing the 5’UTR of the Cln3 mRNA with that of Gal1 mRNA and adjusting Cln3 expression from an inducible promoter to achieve the same mean cell size as WT cells did not cause any statistically significant increase in the spread of cell size, as would be expected if the details of Cln3 synthesis were key elements in cell size regulation. We acknowledge, however, that there may be more subtle effects on the spread in cell size that are unobservable in these measurements on asynchronous populations.

There is clear phenomenological evidence for mechanisms that regulate the distribution of cell sizes: newborn daughters spend longer in G1 than their larger mothers, and very large cells have a smaller exponential growth constant than cells at the median size (48). But what are the molecular mechanisms that produce these correlations? One common theme in the models described above is their focus on the role of single genes in regulating cell size. Although deleting the genes invoked by these hypotheses either increases (*cln3Δ, swi4Δ*) or decreases (*whi3Δ, whi5Δ*) cell size, none of these perturbations produces the dramatic broadening of the cell size distribution expected if a single gene controlled the spread of cell size (15, 16, 49). Our interpretation of our own and earlier work is that an understanding of how cells couple their size to the cell cycle remains elusive. Although the field has identified proteins whose expression alters the average cell size, no model has succeeded in explaining how these proteins collectively prevent the cell size distribution from growing broader over time. We see three classes of model that could close the gap. The first is that there is a single, as yet unidentified, protein that regulates passage through Start, whose abundance or concentration is controlled by cell size in a way that controls the distribution of cell size. The second is that there are multiple proteins that activate passage through Start and multiple proteins that inhibit it, with the synthesis rate of the activators rising with cell size more steeply than that of the inhibitors, so that as cells grow, the activators eventually prevail and drive cells through Start (B. Futcher, personal communication). In this model, size control is a collective exercise, and it would require the removal of many activators or inhibitors to broaden the cell size distribution. Finally, there are likely to be passive mechanisms that regulate cell size in addition to these active mechanisms. As an example, if the ratio between protein synthesis rate and cell volume has an optimal value over a modest range, but then falls in larger or smaller cells, the cells that are too big or too small would replicate more slowly. This would set a limit to the steady state distribution of cell size, even in the absence of active control linking cell cycle progress to cell size. Indeed, recent evidence suggests that the growth of large cells slows because transcription reaches a maximal rate (48, 50). Further measurements of how cell growth rate is affected by cell size will be useful to understand the constraints on the spread of cell size that may be imposed by such a passive size control mechanism in the absence of other active size control strategies (51).

## Methods

See Supplementary Information Section 1 for a full description of the methods used in this text.

### Yeast strains and plasmids

The strains used in this study were congenic with W303 and are listed in full in Table S1. All strains were constructed using standard genetic methods.

### Image processing

We performed brightfield image segmentation and cell tracking using the opensource CellStar algorithm (52). Cell volume was inferred based on this brightfield segmentation by fitting an ellipse to the 2D mask and assuming a 3D prolate spheroid shape. We designed a custom, semi-automated image processing pipeline to incorporate fluorescence data and compile measurements on individual cell cycles. Cell cycle progression was assessed based on Whi5 nuclear localization, in accordance with previous approaches (33). All relevant code is available at https://github.com/AWMurrayLab/image_processing_cellstar_public.git.

### Live cell microscopy

For time-lapse microscopy, cells were loaded into a CellASIC microfluidics flow chamber and imaged using a Nikon Eclipse Ti spinning disc confocal microscope. Growth medium with the appropriate carbon source and galactose concentration was flowed from 2 wells at a pressure of 1PSI using the ONIX microfluidics system. Our single time-point imaging used agar pads with a concentration between 1-2%, made using the appropriate growth medium.

### Cell culture

All experiments were performed in 2X complete synthetic medium (CSM) (53) with various carbon sources. Cells were taken from exponentially growing cultures with a culture density between 1-5 × 10^6^ cells/mL and were grown in the relevant growth medium for a minimum of 20 hours prior to measurement.

## Acknowledgements and Funding Sources

F. B. was supported by the William Georgetti Trust, a Harvard Graduate Merit Award, a Harvard Quantitative Biology Initiative Student Award, The Milton Fund and The Volkswagen Foundation while conducting this research.

A. A. acknowledges the support of the NSF CAREER award number 1752024.

A. W. M. thanks NIH grant RO1-GM43987 and the NSF-Simons Center for Mathematical and Statistical Analysis of Biology at Harvard (#1764269 (NSF) & #594596 (Simons)) for support.

The authors thank Naama Barkai, Ilya Soifer, Angelika Amon, Bruce Futcher and Jan Skotheim for helpful suggestions in writing this manuscript.

F. B. thanks Zachary Niziolek, Jeffery Nelson and the staff at the Harvard Bauer Core Facility for their help with the coulter counter instrument.

