## Supplementary Information for "Cell size regulation in budding yeast does not depend on linear accumulation of Whi5"

#### Section 1: Methods

##### Yeast strains and plasmids

The strains used in this study were congenic with W303 and are listed in full in Table S1. All strains were constructed using standard genetic methods. Fluorescently labeled proteins were constructed to contain a peptide linker between the native protein and the fluorescent label, using a linker sequence that has been previously characterized (1). The fast-maturing mVenNB protein sequence was obtained from as a gift from the Cluzel laboratory (2).

##### Image processing

We performed brightfield image segmentation and cell tracking using the opensource CellStar algorithm (3). Cell volume was inferred based on this brightfield segmentation by fitting an ellipse to the 2D mask. Cell volume was then approximated as that of a prolate spheroid with that given 2D elliptical cross section. We designed a custom, semi-automated image processing pipeline to incorporate fluorescence data and compile measurements on individual cell cycles. Cell cycle progression was assessed based on Whi5 nuclear localization, in accordance with previous approaches (4). Whi5 nuclear localization was detected using an SVM machine learning classifier from the *Scikitlearn* python package, trained on manually annotated data for each individual time-lapse. Since we obtained z-stack time-lapse images, fluorescence signal from the constitutively expressed mCherry protein was used to create 3D masks of each cell. This mask was constructed by thresholding, using a threshold of 3 standard deviations above the average background fluorescence level. As such, these masks did not include vacuoles, since fluorescence was excluded from these organelles. These 3D masks were used to infer average cellular fluorescence as the average of pixel intensities within each mask. All relevant code is available at https://github.com/AWMurrayLab/image_processing_cellstar_public.git.

##### Live cell microscopy

All imaging was performed using a Nikon Eclipse Ti spinning disc confocal microscope, fitted with a Plan Apo 60X/1.40 Oil objective, a Hamamatsu EM-CCD digital camera and a Spectral Borealis Box. EM gain settings were set to within a range of linear signal amplification. All imaging used a spacing of 0.7µm between optical z-sections. For time-lapse microscopy, cells were loaded into a CellASIC microfluidics flow chamber. Growth medium with the appropriate carbon source and galactose concentration was flowed from 2 wells at a pressure of 1PSI using the ONIX microfluidics system. Our single time-point imaging used agar pads with an agar concentration between 1-2%, made using the appropriate growth medium. Cells were taken from exponentially growing cultures and spotted onto pre-made pads, allowed to dry for 10-15 minutes, and imaged using MatTek glass bottom dishes. For our $P_{GAL1}$-*WHI5* and $P_{WHI5}$-*WHI5* strains, we performed time-lapse imaging for 2 experiments with a time-step of 10min, and 1 experiment with a time-step of 12min, to a total of 3 time-lapse experiments per cell type.

##### Cell culture

All experiments were performed in 2X complete synthetic medium (CSM) (5) with various carbon sources. All galactose induction experiments were performed by inoculating cultures in 2X CSM 2% Dextrose, taking cells in log phase between 16-24 hours later, washing in PBS and resuspending in 2X

CSM 3% glycerol for 6 hours to alleviate catabolite repression (6). Cells were then washed in PBS before being grown in 2X CSM with 2% Raffinose and the relevant concentration of Galactose for at least 20 hours prior to measurement. Cells were taken from exponentially growing cultures with a culture density between 0.5-5 x $10^6$ cells/mL.

Coulter counter measurements

To minimize any variability arising from differences in the level of measurement noise, we performed all our Coulter counter measurements with samples containing approximately equal cell concentrations of $4x10^4$ cells/mL. To account for measurement noise, we employed thresholding to exclude datapoints below 10fL, and to exclude the top 2.5% of datapoints. This constrained the effect of extreme outliers on the statistics describing the spread in cell size. These thresholds did not significantly alter the relative standard deviation or CV measurements compared between different samples, but generated statistics that represented the bulk of the size distribution. We were able to reproduce these statistics computationally by using our time-lapse measurements of the spread in cell size at birth, and assuming exponential growth in an age-structured population.

Simulations of population growth

The simulations in Figure S1 were performed using a model in which cells pass through Start stochastically at a rate that depends inversely on the Whi5 concentration, i.e.

$$\mu([W]) = \frac{k}{[W]^n},$$

where $k$ is a rate constant, $[W]$ is the concentration of Whi5, and $n$ is an exponent (taken for the simulations in Figure S1 to be $n=2$). Here $\mu$ represents the rate of passage through Start per unit time for a cell with a given concentration of Whi5, with division occurring after a constant time interval post-Start. For simplicity we took the division asymmetry between mother and daughter cells to be constant, with a constant ratio $r$ between daughter cell volume and mother cell volume at division. This model was constructed to represent one realization of the general stochastic rate model described in (7). We tested a model with this rate of passage through Start for two different synthesis profiles of Whi5. The first is described further in (8) and corresponds to one in which a constant amount $\Delta$ of Whi5 is produced between Start and division, which is then diluted during the subsequent G1 phase. This represents the "WT" case in Figure S1 due to its consistency with previous studies of the production rate of Whi5 (7). We note that this first model is capable of generating size control, with the average cell size, standard deviation in cell size and CV remaining robustly regulated throughout our simulations (see Figure S1).

The second model for Whi5 synthesis is one in which Whi5 is produced at a rate proportional to volume:

$$\frac{dW}{dt} = k_w V(t)$$

Where $k_w$ is the production rate of Whi5 for a given cell volume $V$, and W is the abundance of Whi5. This leads to a steady state concentration of Whi5 $[W]^* = \frac{k_w}{\lambda}$, where $\lambda$ is the exponential volume growth rate. Within the second model, the attainment of a steady state Whi5 concentration leads cells to proceed through Start with a constant rate $\mu = \frac{k\lambda^n}{k_w^n}$, independent of their cell volume. Passage times through Start will then follow an exponential distribution with rate parameter $\mu$. The average relative growth acquired during the G1 phase is then $E\left(\frac{V_s}{V_b}\right) = E\left(e^{\lambda t}\right) = \frac{\mu}{\mu-\lambda}$. We therefore simulated cell growth, varying the synthesis rate $k_w$ to test different regimes of this model. We tested three cases: high $k_w = 3.0$ to generate excess growth in G1 with a high concentration of Whi5, small $k_w = 0.1$ to generate only small amounts of growth in G1, with a low concentration of Whi5, and an intermediate synthesis rate $k_w = 0.7$. This third case was selected to generate an average cell size which matched that of "WT" cells simulated with a volume independent production of Whi5 each cell cycle (see Figure S1). For concreteness, we performed our simulations for the case *n=2, λ=1, k=2, r=0.5*, but our conclusions about the inability of the second model to produce size control do not rely on a specific choice of these parameters. Over the range of parameters we tested, this second model is incapable of generating size control, with the average cell size and standard deviation in cell size either arbitrarily increasing or decreasing depending on the rate of Whi5 production. In either case, this second model gives rise to an unconstrained increase in the CV over successive generations (Figure S1). This result is to be expected, since any model in which the rate of passage through Start $\mu$ is determined by the Whi5 concentration will display a constant rate $\mu$ if the Whi5 concentration reaches a steady state value. A constant rate of passage through Start will not provide feedback towards a mean cell size, and errors in interdivision times are therefore expected to accumulate over successive cell cycles, similar to the geometric random walk predicted for symmetrically dividing cells (9).

Our simulations were performed using a discretized time approach, in which the probability of cells passing through Start in a given time interval *dt* was taken to be *μdt*. To simulate population growth for many generations we implemented a Moran process; for populations that exceeded a size of 10,000 cells, each cell division event was accompanied by the selection and removal of a cell at random. The code for our simulations is available online at https://github.com/AWMurrayLab/growth_rate_simulations_public.git, and has been described in greater detail previously (8).

#### Section 2: Supplementary Tables

**Table S1: *S. cerevisiae* strains and plasmids used in this study.**

| *S. cerevisiae* strain label | Genotype | Source | Figures this strain was used in |
|---|---|---|---|
| yJHK85 | *MATa; bud4-W303; BUD4 (S288C); can1-100; his3-11,15; ura3Δ* | Gift from John Koschwanez (Murray Lab) | NA |
| yFB29 | *yJHK85; bud4-W303; BUD4 (S288C); can1-100; his3-11,15; ura3Δ; gal3Δ:PrACT1-GAL3-TrGAL3; gal1Δ,gal10Δ:His3MX6; whi5Δ:KanMX-*$P_{GAL1}$*-WHI5-linker-mVenNB* | This study | S2, S3 |
| yFB30 | *yJHK85; bud4-W303; BUD4 (S288C); can1-100; his3-11,15; ura3Δ; gal3Δ:PrACT1-GAL3-TrGAL3; gal1Δ,gal10Δ:His3MX6; whi5Δ:KanMX-*$P_{GAL1}$*-WHI5-linker-mVenNB; bck2Δ:NatMX* | This study | S2, S3 |
| yFB43 | *yJHK85; bud4-W303; BUD4 (S288C); can1-100; his3-11,15; ura3Δ; gal3Δ:PrACT1-GAL3-TrGAL3; gal1Δ,gal10Δ:His3MX6; whi5Δ:WHI5-linker-mVenNB-KanMX* | This study | S2, S3 |
| yFB46 | *yJHK85; bud4-W303; BUD4 (S288C); can1-100; his3-11,15; ura3Δ; gal3Δ:PACT1-GAL3-TrGAL3; gal1Δ,gal10Δ:His3MX6; whi5Δ:WHI5-linker-mVenNB-KanMX; bck2Δ:NatMX* | This study | S2, S3 |
| yFB78 | *yJHK85; bud4-W303; BUD4 (S288C); can1-100; his3-11,15; ura3Δ; gal3Δ:*$P_{ACT1}$*-GAL3-TrGAL3; gal1Δ,gal10Δ:His3MX6; whi5Δ:KanMX-*$P_{GAL1}$*-WHI5-linker-mVenNB; act1Δ: mCherry-*$T_{ADH1}$*-HPHMX4-*$P_{ACT1}$*-ACT1* | This study | 2, 3, 4, S2, S3, S4, S5, S6, S7, S8, S9 |
| yFB79 | *yJHK85; bud4-W303; BUD4 (S288C); can1-100; his3-11,15; ura3Δ; gal3Δ:PrACT1-GAL3-TrGAL3; gal1Δ,gal10Δ:His3MX6; whi5Δ:WHI5-linker-mVenNB-KanMX; act1Δ: mCherry-*$T_{ADH1}$*-HPHMX4-*$P_{ACT1}$*-ACT1* | This study | 2, 3, 4, S2, S3, S4, S5, S6, S7, S8, S9 |
| yFB86 | *yJHK85; bud4-W303; BUD4 (S288C); can1-100; his3-11,15; ura3Δ; gal3::PrACT1-GAL3-TrGAL3; gal1Δ, gal10Δ::His3MX6; cln3Δ::KANMX-*$P_{GAL1}$*-CLN3* | This study | S10 |
| Plasmid label | Genotype | Source | |
| pFB9 | *pUC19* with the following *WHI5* cassette inserted at the Sma1 cut site: $P_{WHI5}$*-WHI5-linker-mVenNB-*$T_{ADH1}$*-*$P_{TEF}$*-KanMX-*$T_{TEF}$*-*$T_{WHI5}$ | This study | |

| Table S2: Comparison of division times measured with those of Di Talia *et al*. (4) | | |
|---|---|---|
| **Measurement type** | **Value** | **Source** |
| $T_{G1}$ in Daughters | 94±3 (108) | Di Talia et al. |
| $T_{G1}$ in Mothers | 8±1 (178) | Di Talia et al. |
| Total cycle Daughter | 219±3 (54) | Di Talia et al. |
| Total cycle Mother | 133±2 (84) | Di Talia et al. |
| $T_{G1}$ in Daughters | 123±2 (853) | This study |
| $T_{G1}$ in Mothers | 40±1 (1581) | This study |
| Total cycle Daughter | 224±2 (853) | This study |
| Total cycle Mother | 137±1 (1581) | This study |

**Table S2:** Comparison of division time statistics measured in this study on our $P_{WHI5}$-*WHI5* haploid cells grown in 2% Raffinose with those acquired separately on WT haploid cells grown in glycerol/ethanol as a carbon source. Values in rows 1-4 are quoted from Tables S5 and S6 in Di Talia *et al.* (4) and are re-labeled here for consistency with our measurements of G1 duration as the period from Whi5 nuclear entry to Whi5 nuclear exit. Growth in Raffinose displays an extended G1 phase, but a similar overall cell cycle duration. The table shows the mean +/- the standard error of the mean in minutes, with the number of cells observed reported in parentheses.

### Section 3: Supplementary Figures

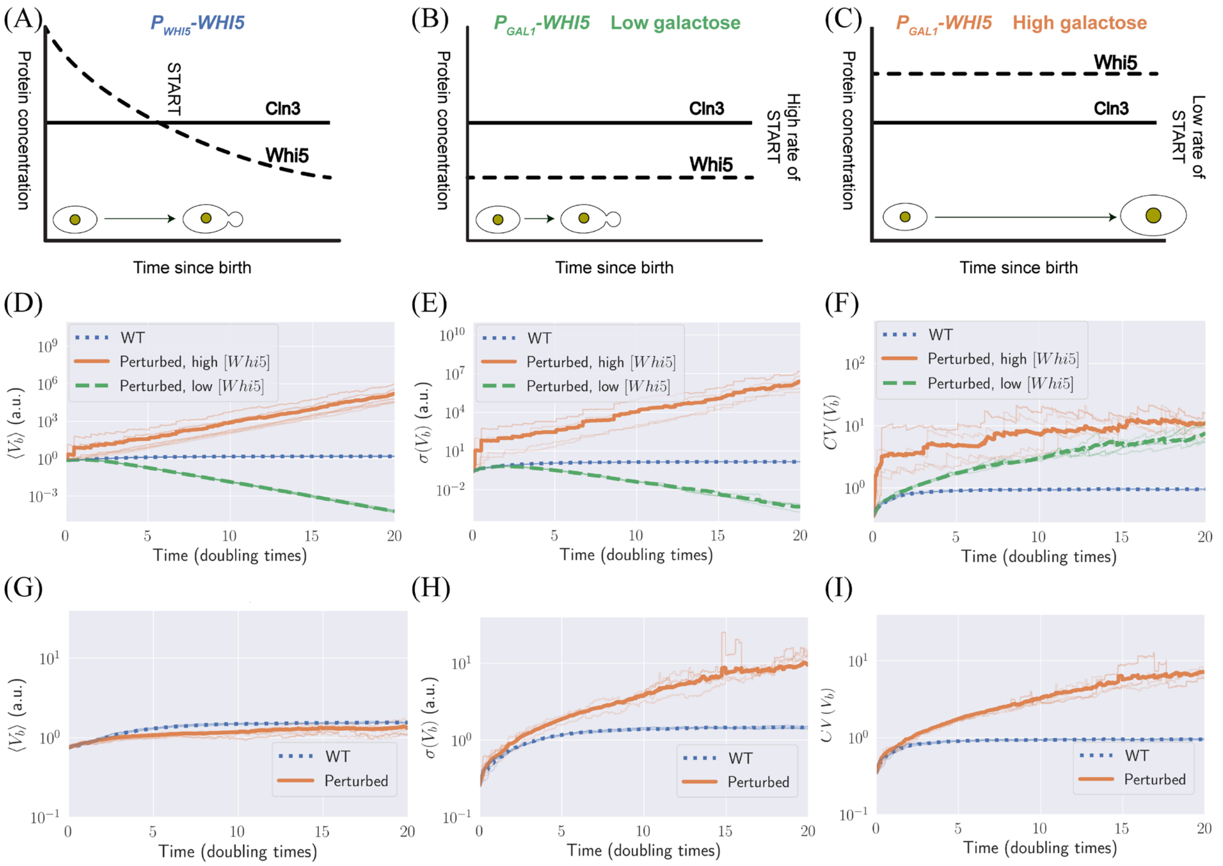


**Figure S1:** (A-C) A schematic representation of Whi5 and Cln3 concentrations throughout G1 in (A) WT cells, showing Whi5 dilution with cell growth and (B-C) two different, constant concentrations of Whi5 produced by growing cells in low (B) or high (C) concentration of galactose. (D-I) Simulations of a noisy rate model for passage through Start. We simulated the growth of populations of cells in which Whi5 was produced independently of cell volume ("WT" cells in blue), or at a rate which scaled with cell volume ("Perturbed" cells in green or orange). The regular discontinuities correspond to a numerical artefact of regularly sampling a subset of cells to limit the total population size. This is necessary to avoid the population size growing excessively large and exceeding computer memory limitations. Thick lines correspond to the average over 10 simulated repeats, and thin lines correspond to individual repeats. (D-F) Average cell volume at birth, standard deviation in volume at birth, and CV in volume at birth plotted against the number of population doubling times. Cells with high Whi5 concentration (orange) increase arbitrarily in average cell size and standard deviation in cell size. Cells with low Whi5 concentration (green) decrease arbitrarily in average cell size and standard deviation in cell size. In contrast, WT cells (blue) show little variability in average cell size and spread in cell size over the range of these simulations. (G-I) The orange curve shows simulated cell growth when the steady state Whi5 concentration is tuned to generate an average cell size matching that of WT cells. In this case we still observe an increase in the standard deviation and CV in cell volume (measured at birth) which is unconstrained for the duration of our simulations relative to WT cells which produce a constant amount of Whi5 with each cell cycle. All other simulation parameters remain fixed between WT cells and cells in which Whi5 production is proportional to cell volume. Details for simulations in (D-I) are contained in Supplementary Information Section 1.

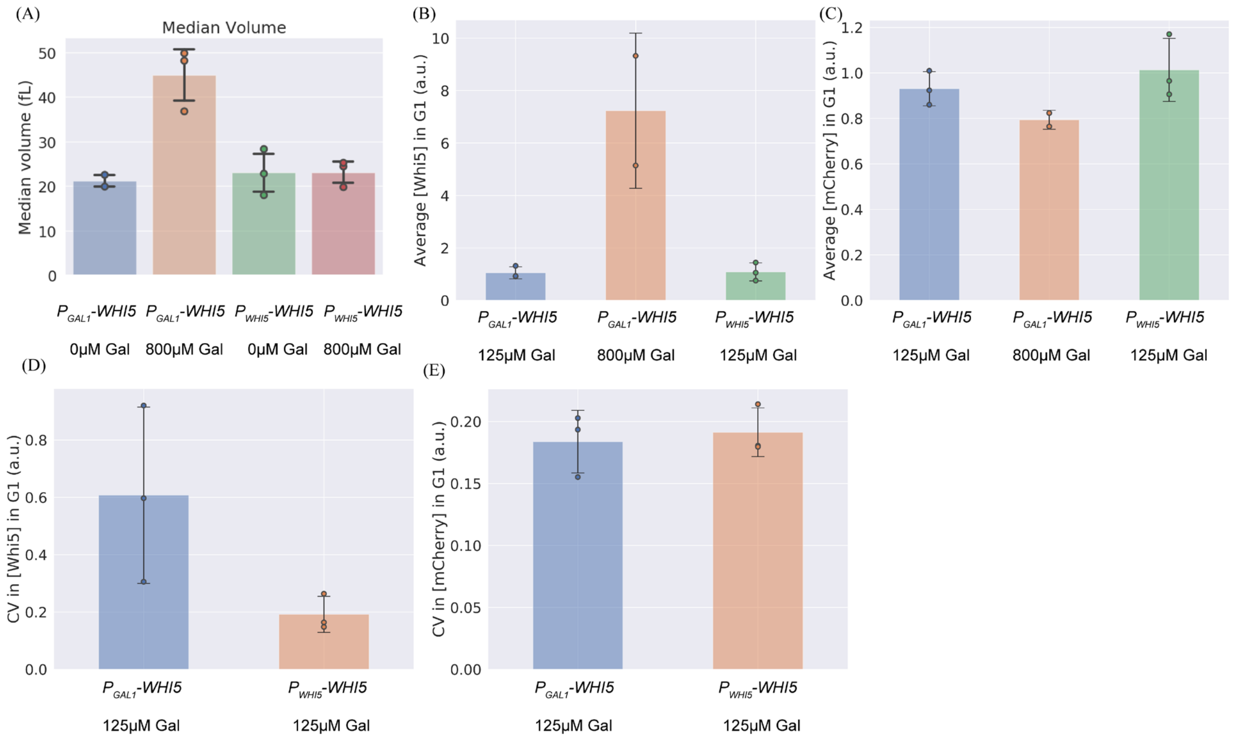


**Figure S2:** Overexpressing Whi5 from the *GAL1* promoter substantially increases cell size, concomitant with an increase in Whi5 concentration. (A) Measurements of median cell size for fluorescently labeled Whi5 in cells expressing *WHI5* from $P_{GAL1}$ in 0μM and 800μM Galactose, obtained using a Coulter counter on asynchronous populations of log phase cells. We note that $P_{GAL1}$-*WHI5* cells grown without galactose did not show a substantial decrease in cell size below that of our $P_{WHI5}$-*WHI5* cells, which may indicate a low basal level of expression from our galactose inducible system in these growth conditions. $P_{WHI5}$-*WHI5* cells show no dependence of cell size on galactose concentration in this condition. (B-C) Fluorescence concentration measurements for populations of G1 cells, obtained via single time-point microscopy. (B) Whi5 concentration shows a marked increase in $P_{GAL1}$-*WHI5* cells grown with 800μM Galactose. (C) mCherry concentration, expressed from the *ACT1* promoter, remains relatively constant across different galactose concentrations. (D-E) Measurements of the CV in Whi5 and mCherry concentration measured by time-point microscopy, showing an increased CV in $P_{GAL1}$-*WHI5* expression relative to $P_{WHI5}$-*WHI5* cells. Error bars represent standard deviation calculated over at least two experimental replicates per condition for the relevant statistic. Dots correspond to values for individual biological replicates.

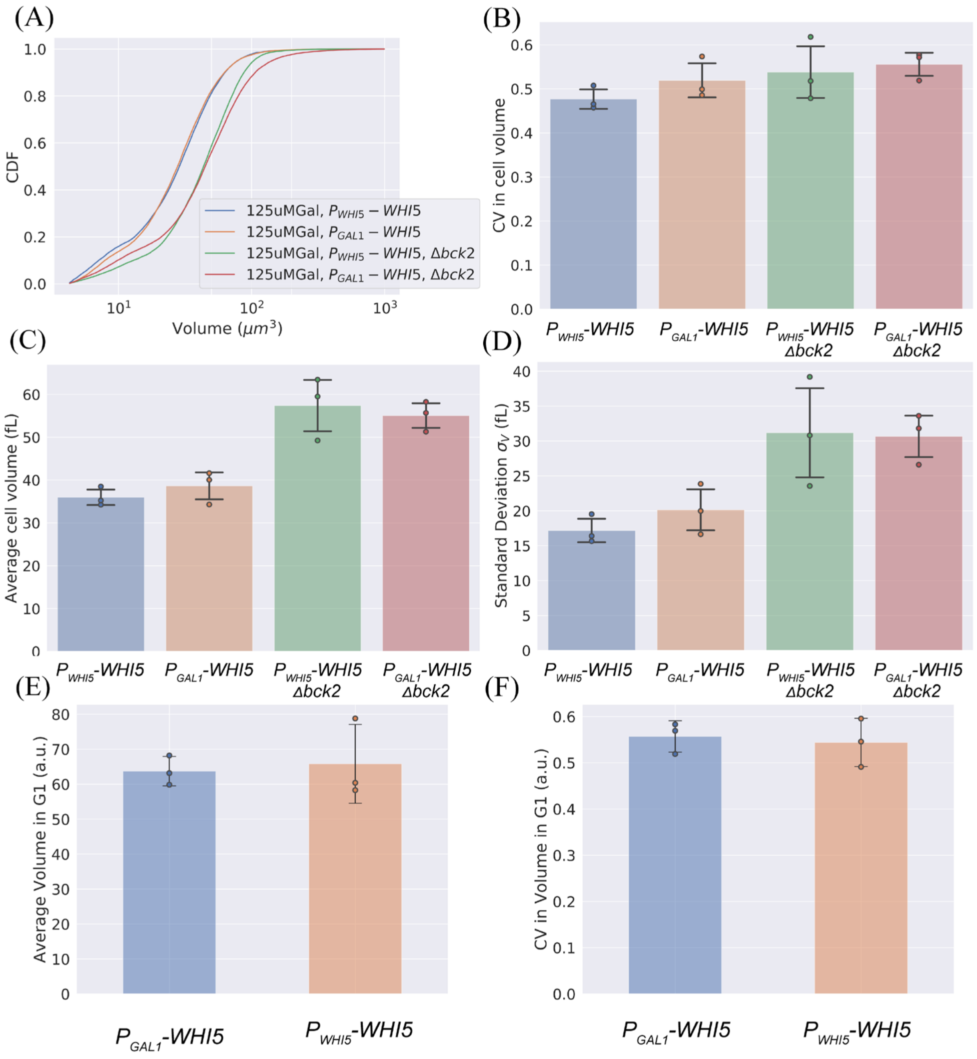


**Figure S3:** The spread in cell size measured in asynchronous populations is statistically indistinguishable whether Whi5 is driven by its own promoter or the *GAL1* promoter. We tuned the expression of the *GAL1* promoter, using 125μM galactose, to give a level of Whi5 expression that gave the same average cell size as the WT ($P_{WHI5}$-*WHI5*). (A-D) Size statistics measured by Coulter Counter. (E-F) Size statistics for G1 cells measured by single time-point microscopy. (A) The cumulative distribution function of the size distribution for $P_{GAL1}$-*WHI5* cells and $P_{WHI5}$-*WHI5* cells in *BCK2* and *bck2Δ* backgrounds. This data is representative of 3 experimental replicates. (B-D) Statistics (CV, average cell size, and standard deviation in cell size) of the size distributions from (A). Error bars are the standard deviation of the statistic measured across at least 3 experimental replicates. The spread of the cell size distribution is identical whether Whi5 is expressed from its own promoter or from $P_{GAL1}$, in both *BCK2* and *bck2Δ* cells, inconsistent with the predictions of the inhibitor dilution model. (E) Average volume in G1 for an asynchronous population, measured by time-point microscopy. (F) CV in volume in G1 for an asynchronous population, measured by time-point microscopy. The values reported in (E-F) neglected cells with volume above 200fL, to avoid skewing the cell size distribution with large cells that have arrested in G1 (Neurohr et al., 2019). The effect of this thresholding decreases the CV for both cell types but does not alter the size of their CVs relative to each other. Performing a Welch's t-test across experimental replicates shows no statistically significant differences when comparing *$P_{GAL1}$-WHI5* vs. *$P_{WHI5}$-WHI5* cells. We note that the G1 CVs observed in the asynchronous G1 cell time-point data can be recapitulated computationally using populations seeded with the CVs measured in cell volume at birth, providing an internal consistency check between these two distinct approaches. Dots correspond to values for individual biological replicates.

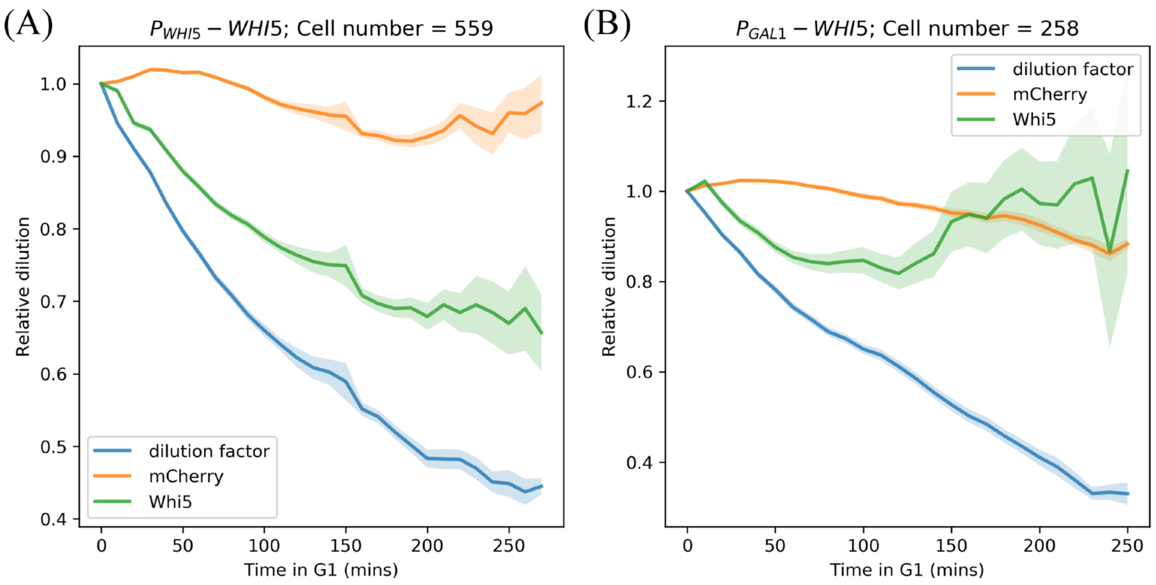


**Figure S4:** Relative protein concentration throughout G1 for daughter cells, measured via fluorescence intensity normalized to intensity at birth. Shaded bars correspond to standard error of the mean across the population of cells at each timepoint. The blue dilution factor shows the expected relative dilution of a given fixed protein abundance based on the incremental volume added during G1 growth (i.e. $V_b/V(t)$). (A) $P_{WHI5}$-*WHI5* cells show decreasing fluorescence intensity throughout during G1, in contrast to a relatively constant fluorescence intensity generated by $P_{ACT1}$ driving *mCherry* expression. (B) $P_{GAL1}$-*WHI5* cells do not replicate the dilution behavior seen in $P_{WHI5}$-*WHI5* cells. Data is compiled for two experiments, which used the same time-steps, 10 minutes, for imaging and results are qualitatively similar to a third experiment which used a 12-minute time-step.

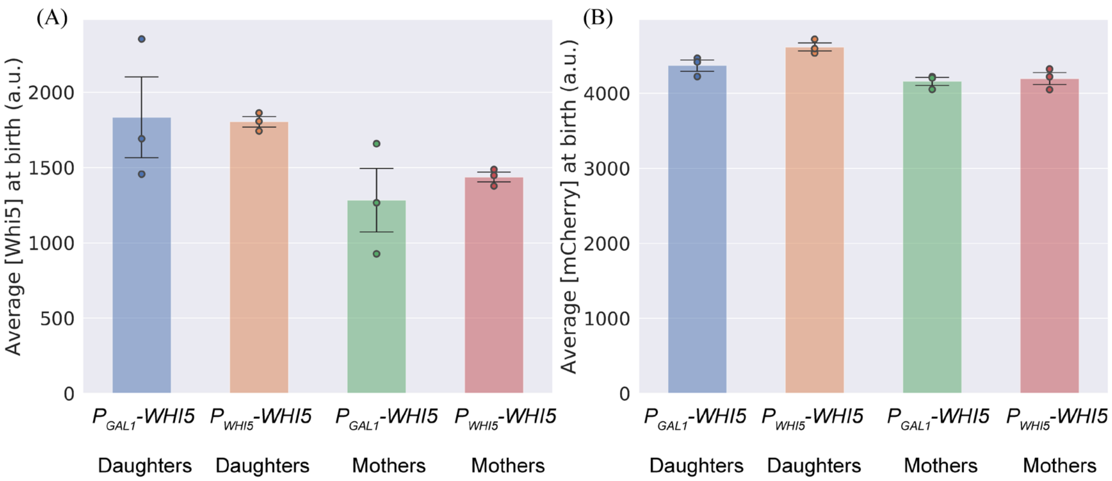


**Figure S5:** Measurements of Whi5 and mCherry concentration at cell birth, measured via time-lapse microscopy on cells grown with 125µM Galactose. (A) Average Whi5 concentration at birth, grouped by cell type. (B) Average mCherry concentration at birth, grouped by cell type. Error bars correspond to the standard deviation measured over multiple biological replicates. Performing a Welch's t-test across biological replicates shows no statistically significant differences when comparing *$P_{GAL1}$-WHI5 vs. $P_{WHI5}$-WHI5* cells for daughters and mothers separately. Dots correspond to values for individual biological replicates.

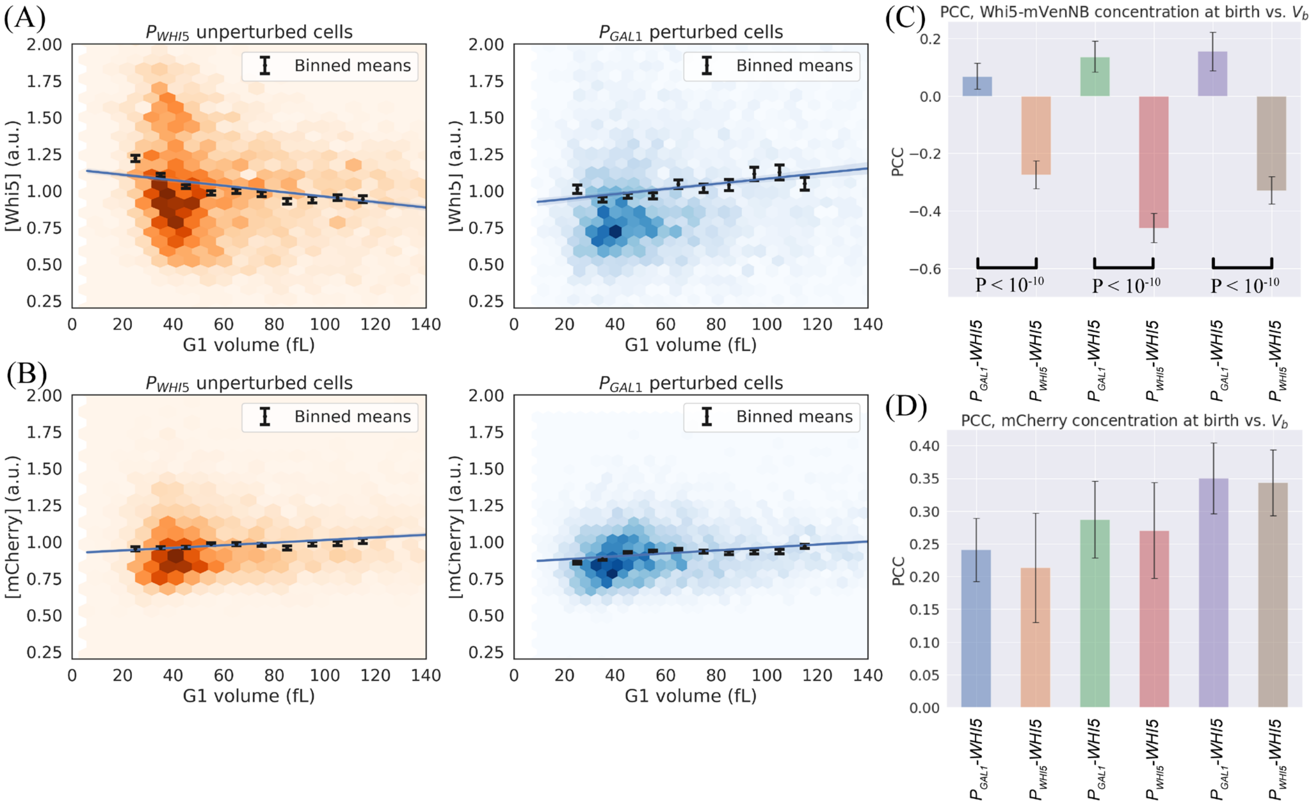


**Figure S6:** (A-B) Plots of protein concentration vs. volume for G1 cells, measured via single timepoint microscopy on cells grown with 125uM Galactose. Colored hexagons represent a 2D histogram of datapoints, with darker hexagons showing increased local density of data points. Black lines correspond to averages of the same data binned with respect to $V_b$, with error bars showing the standard error of the mean. Blue lines correspond to linear regression fits, with 95% confidence intervals. Data is compiled from three experiments for each cell type. (A) [Whi5] vs. volume in G1 shows a consistent negative correlation for $P_{WHI5}$-*WHI5* cells, in contrast to a positive correlation observed for $P_{GAL1}$-*WHI5* cells. (B) No difference between cell types is observed for the correlation between [mCherry] and cell volume. (C-D) Pearson correlation coefficients plotted in pairs for the three biological replicates plotted in (A-B). Black lines correspond to statistically significant differences with P values less than 0.05 quoted, calculated using a Fisher's z-transformation on both datasets. (C) Statistically significant differences are consistently observed by comparing $P_{WHI5}$-*WHI5* and $P_{GAL1}$-*WHI5* cell types within each biological replicate for the Whi5 signal. (D) No statistically significant differences are observed by comparing $P_{WHI5}$-*WHI5* and $P_{GAL1}$-*WHI5* cell types within each biological replicate for the mCherry signal.

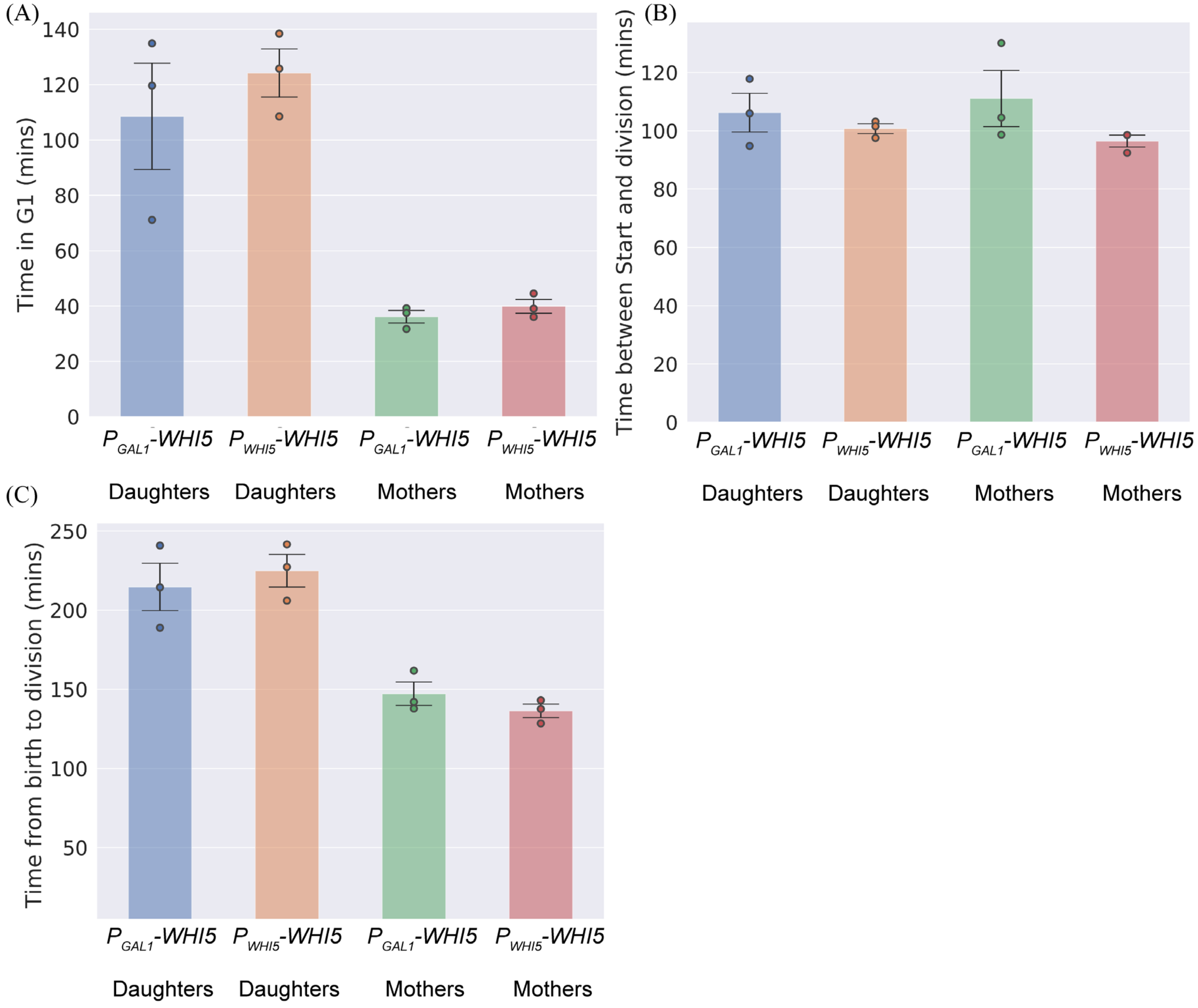


**Figure S7:** Cell cycle timing is unperturbed in $P_{GAL1}$-*WHI5* and $P_{WHI5}$-*WHI5* cell types. Plots of the cell cycle timing in minutes grouped by cell type, measured by time-lapse microscopy on cells grown with 125µM Galactose. (A) Average time in the G1 phase. (B) Average budded duration. (C) Average division time. Performing a Welch's t-test across biological replicates shows no statistically significant differences when comparing daughters with daughters and mothers with mothers for $P_{GAL1}$-*WHI5 vs.* $P_{WHI5}$-*WHI5* cells. Dots correspond to values for individual biological replicates.

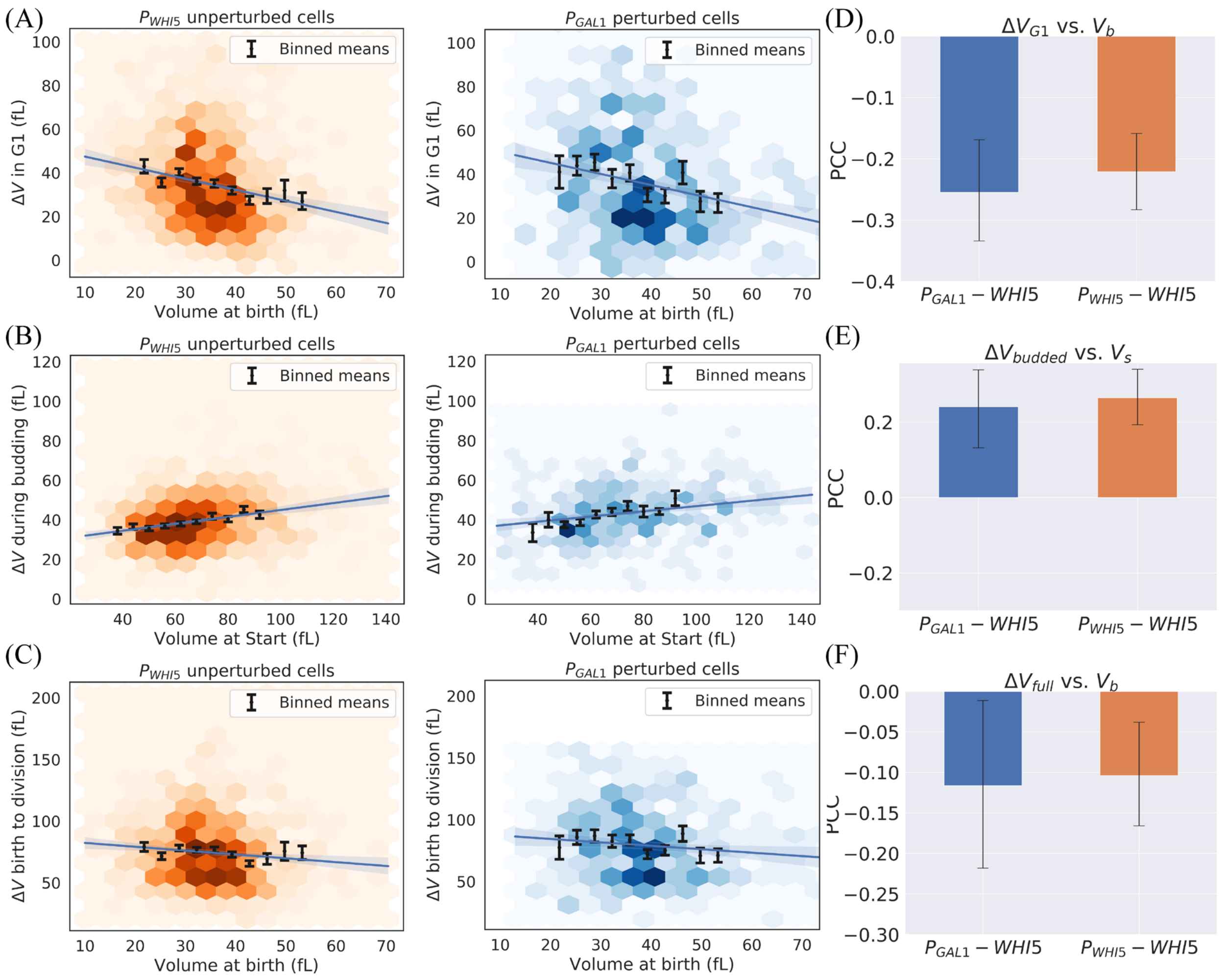


**Figure S8:** Plots showing the dependence of volume added between birth and Start, between Start and division, and over the entire cell cycle vs. cell volume at birth and at Start. Plots are shown for $P_{GAL1}$-*WHI5* (blue) and $P_{WHI5}$-*WHI5* (orange) cell types. See Figure S6 for details on plotting features for (A-C). (A) Plots showing volume added during G1 vs. cell volume at birth. (B) Plots showing volume added between Start and division vs. cell volume at Start. (C) Plots showing volume added over the full cell cycle vs. cell volume at birth. (E-F) PCC measurements for the data sets plotted in (A-C). No statistically significant differences are observed between cell types, as indicated by performing a Fisher's z-transformation on both datasets.

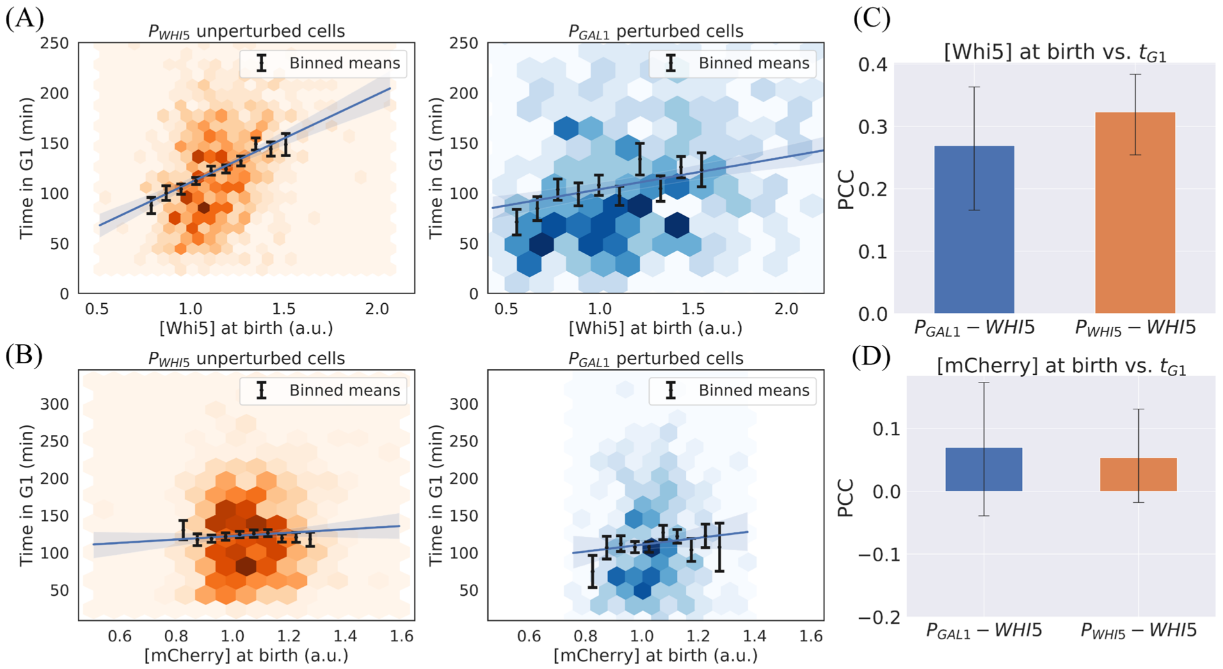


**Figure S9:** Plots of the dependence of G1 duration on the concentration at birth of (A) Whi5 and (B) mCherry (presented for completeness), measured via time-lapse microscopy, using fluorescence intensity as a proxy for protein concentration. See Figure S6 for details on plotting features for (A-B). Data is compiled from three experiments for each cell type. (A) G1 duration maintains a positive correlation with Whi5 concentration at birth in $P_{GAL1}$-*WHI5* and $P_{WHI5}$-*WHI5* cell types. (B) G1 duration shows a weak positive correlation with mCherry concentration at birth. (C-D) PCC calculations for the data sets plotted in (A-B). No statistically significant difference is observed between cell types, as indicated by performing a Fisher's z-transformation on both datasets. We note that although based on a PCC comparison the correlations presented in (A) do not show any statistically significant difference, the slope of a $P_{GAL1}$-*WHI5* linear regression is less steep than that observed for the $P_{WHI5}$-*WHI5* strain ($P<10^{-4}$). In fact, a weakened dependence of time in G1 on Whi5 concentration at birth might be expected for this strain, since Whi5 concentration at birth is no longer negatively correlated with cell volume in $P_{GAL1}$-*WHI5* cells for this condition.

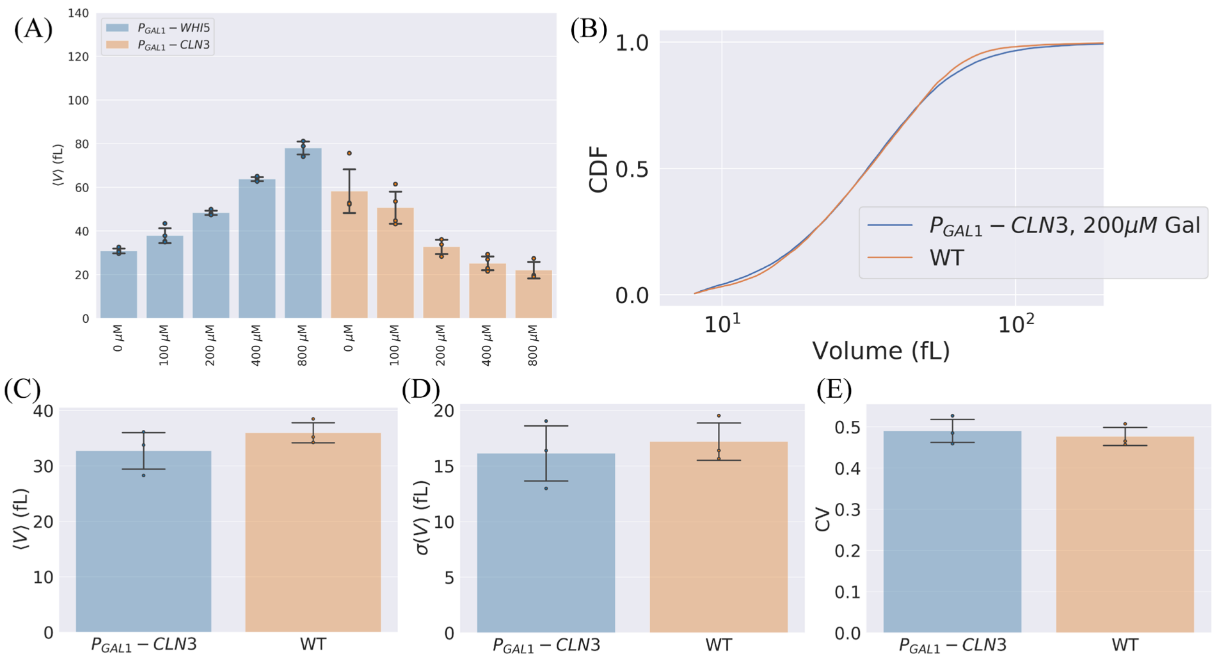


**Figure S10:** Size statistics of *P<sub></sub>*$P_{GAL1}$-*WHI5*, $P_{GAL1}$-*CLN3* and WT strains. Data acquired on asynchronous populations by Coulter counter. (A) Average size measured at various galactose concentrations (0µM through to 800µM) for $P_{GAL1}$-*WHI5* (blue) and $P_{GAL1}$-*CLN3* (orange) strains. (B) Cumulative distribution functions (CDFs) plotted for WT cells (orange) and $P_{GAL1}$-*CLN3* cells (blue) where the galactose concentration has been tuned to 200µM, generating cells with the same average cell size as WT cells. Data is plotted only for one repeat but is representative of multiple biological replicates. (C, D, E) Cell volume statistics for WT cells and $P_{GAL1}$-*CLN3* cells grown with 200µM galactose. (C) Average cell volume. (D) Standard deviation σ in cell volume. (E) CV in cell volume. Bar plots show the average over 3 biological replicates, while dots represent individual biological replicates. No statistically significant differences are observed between the two cell types in C-E (Welch's T-test).
